# The effect of perinatal brain injury on dopaminergic function and hippocampal volume in adult life

**DOI:** 10.1101/128710

**Authors:** Sean Froudist-Walsh, Michael A.P. Bloomfield, Mattia Veronese, Jasmin Kroll, Vyacheslav Karolis, Sameer Jauhar, Ilaria Bonoldi, Philip K. McGuire, Shitij Kapur, Robin M. Murray, Chiara Nosarti, Oliver D. Howes

## Abstract

**Background:** Very preterm birth (<32 weeks of gestation) is associated with long-lasting brain alterations and an increased risk of psychiatric disorders associated with dopaminergic abnormalities. Preclinical studies have shown perinatal brain injuries, including hippocampal lesions, cause lasting changes in dopamine function, but it is not known if this occurs in humans. The purpose of this study was to determine whether very preterm birth and perinatal brain injury were associated with altered dopamine synthesis and reduced hippocampal volume in humans in adulthood.

**Methods:** We compared adults who were born very preterm with associated perinatal brain injury to adults born very preterm without perinatal brain injury, and age-matched controls born at full term using [18F]-DOPA PET and structural MRI imaging.

**Results:** Dopamine synthesis capacity was significantly reduced in the perinatal brain injury group relative to both the group born very preterm without brain injury (Cohen’s d=1.36, p=0.02) and the control group (Cohen’s *d*=1.07, p=0.01). Hippocampal volume was reduced in the perinatal brain injury group relative to controls (Cohen’s *d* = 1.17, p = 0.01). There was a significant correlation between hippocampal volume and striatal dopamine synthesis capacity (r = 0.344, p= 0.03).

**Conclusions:** Perinatal brain injury, but not very preterm birth without macroscopic brain injury, is associated with persistent alterations in dopaminergic function and reductions in hippocampal volume. This is the first evidence in humans linking neonatal hippocampal injury to adult dopamine dysfunction, and has implications for understanding the mechanism underlying cognitive impairments and neuropsychiatric disorders following very preterm birth.

## Introduction

More than 10% of babies born in the USA are born preterm (born before 37 weeks of gestation), and about 2% are born very preterm (VPT, before 32 weeks of gestation) (1). Premature birth is a risk factor for cognitive impairment (2) and a number of psychiatric disorders, including schizophrenia and affective disorders (3).

The second and third trimesters of gestation are critical periods for neurodevelopment, particularly for axon and synapse formation, glial proliferation and the development of neurotransmitter systems including the dopaminergic system (4). Thus VPT birth occurs during a critical time for the development of a number of neural systems, when the brain is particularly susceptible to exogenous and endogenous insults (5). VPT babies are at risk of sustaining a variety of perinatal brain injuries, including periventricular haemorrhage, ventricular dilatation and periventricular leukomalacia that are often associated with hypoxic-ischaemic events (6).

The sequelae of VPT birth include long-lasting and widespread structural brain alterations, with reduced hippocampal volume being one of the most consistent findings (7). There is substantial evidence from animal models that perinatal brain injury due to hippocampal lesions (8) or obstetric complications (9) can lead to long-term alterations in the dopamine system, which remain evident in adulthood. Some neonatal insult models have found attenuated striatal dopamine release (10, 11). Recent post-mortem evidence in humans suggests that prolonged early hypoxia can lead to reduced expression of tyrosine hydroxylase, the rate-limiting enzyme for dopamine synthesis (12). In contrast, other animal models of neonatal insults have provided evidence for striatal hyperdopaminergia, with subtle neonatal experimental manipulations sometimes having complex effects on dopaminergic function tested later in life (9). This may mirror the increased dopamine synthesis and release seen in human schizophrenia (13), a condition that has long been associated with obstetric complications (14).

However it is not known if perinatal brain injury is associated with dopaminergic alterations in adulthood in humans, or how this relates to specific brain alterations. We aimed to disentangle the preclinical, post-mortem and indirect clinical evidence regarding the effects of early brain insults on later dopamine function by directly comparing two contrasting hypotheses, namely that early brain injury leads to hyper-, or alternatively hypo-dopaminergia in the striatum. Moreover, in view of the preclinical findings showing that perinatal hippocampal lesions can lead to lasting alterations to the dopamine system (10), and the vulnerability of the hippocampus to perinatal brain injury (15), we hypothesised that hippocampal volume and striatal dopaminergic function would be related.

We set out to test these hypotheses by studying adults who were born VPT with evidence of macroscopic perinatal brain injury who have been followed longitudinally for their entire lives, and compared them to two control groups, one group of individuals born VPT without evidence of macroscopic perinatal brain injury, and a group of controls without a history of VPT birth or perinatal brain injury.

## Method

### Participants

We assessed a group of individuals born VPT who were admitted to the Neonatal Unit of University College Hospital, London in 1979-1985. These individuals were enrolled in a longitudinal study and have been studied periodically for their entire lives.

Macroscopic perinatal brain injury was qualitatively assessed in all participants born VPT and diagnosis of perinatal brain injury was made after consensus between at least two neuroradiologists with a special interest in neonatology. Hemorrhage into the germinal matrix, and those extending to the lateral ventricles or brain parenchyma was labeled as periventricular hemorrhage (16), with the grade defined according to the criteria described by Papile and colleagues (17). Ventricular dilatation was defined as visible dilatation of the lateral ventricles with cerebrospinal fluid while being insufficient to meet the criteria for hydrocephalus. We compared the perinatal brain injury group to: 1) a group of VPT individuals who were similarly assessed at birth but not diagnosed as having perinatal brain injury (to control for the effects of preterm birth) and 2) healthy controls without a history of perinatal brain injury or preterm birth (control group).

Participants who gave consent at previous study time-points to be contacted regarding the study were recruited using the contact details provided previously, and control participants were recruited via advertisements in the local community. Exclusion criteria for all groups were history of post-natal head injury, neurological condition (including stroke, meningitis, multiple sclerosis, and epilepsy) or significant physical illness (such as endocrine or metabolic disorder requiring treatment), substance dependence or abuse, psychotic disorder, current antipsychotic use, and pregnancy. The study was undertaken with the understanding and written informed consent of each subject, with the approval of the London Bentham Research Ethics Committee, and in compliance with national legislation and the Code of Ethical Principles for Medical Research Involving Human Subjects of the World Medical Association (Declaration of Helsinki). Birth weight was recorded for all VPT participants and socio-economic status measured in all subjects using the Standard Occupational Classification (18).

### PET data acquisition

In adulthood, all participants underwent a 3,4-dihydroxy-6-[18F]-fluoro-/-phenylalanine ([18F]-DOPA) scan in a Biograph 6 PET/CT scanner with Truepoint gantry (SIEMENS, Knoxville, TN). Subjects were asked to fast from midnight and abstain from smoking tobacco and consuming food and liquids (except for buttered toast and water) from midnight before the day of imaging to ensure there were no group differences in amino acid consumption prior to the scan. On the day of the PET scan, a negative urinary drug screen was required and a negative pregnancy test was required in all female subjects. Subjects received carbidopa 150 mg and entacapone 400 mg orally 1 hour before imaging to reduce the formation of radiolabeled [18F]-DOPA metabolites (19, 20). Head position was marked and monitored via laser crosshairs and a camera, and minimized using a head-strap. A transmission CT scan was performed before radiotracer injection for attenuation and scatter correction. Approximately 150 MBq of [18F]-DOPA was administered by bolus intravenous injection 30 seconds after the start of PET imaging. We acquired emission data in list mode for 95 minutes, rebinned into 26 time frames (30-second background frame, four 60-second frames, three 120-second frames, three 180-second frames, and fifteen 300-second frames).

### MRI data acquisition

On a separate day an MRI scan was performed on a 3 Tesla GE Signa MR scanner (GE Healthcare). T1-weighted images were acquired (TR/TE/TI: 7.1/2.8/450ms, matrix: 256x256), allowing for 196 slices with no gap and an isotropic resolution of 1.1x1.1x1.1mm^3^.

## Image preprocessing

To correct for head movement, nonattenuation-corrected dynamic images were denoised using a level 2, order 64 Battle-Lemarie wavelet filter (21), and individual frames were realigned to a single frame acquired 10 minutes after the [18F]-DOPA injection using a mutual information algorithm (22).

Transformation parameters were then applied to the corresponding attenuation-corrected frames, and the realigned frames were combined to create a movement-corrected dynamic image (from 6 to 95 minutes following [18F]-DOPA administration) for analysis.

Automatic reconstruction of the hippocampus, caudate nucleus, putamen, nucleus accumbens and cerebellum was performed in the native space of each of the participants with MRI data, allowing for both individual masks and regional volume information extraction, using FreeSurfer version 5.1 (23). The primary striatal region of interest was the whole striatum (nucleus accumbens, caudate and putamen combined) but we also report the sub-regions separately to determine if there were sub-regional variations.

A linear transformation was created between each participant’s T1-weighted structural scan and their individual PET image using FSL FLIRT (24). This transformation was then applied to each of the previously specified regions of interest in order to obtain individually defined masks of the striatum on the PET scan. Intra-subject registration is generally more accurate than between-subject registration, as there is no between-subject anatomical variability to take into account.

In addition to the two participants (one perinatal brain injury, one very preterm no diagnosed injury) excluded from the PET study, four further participants (three perinatal brain injury, one control) were not included in the MRI study due to contraindications to scanning. In order allow for the inclusion of these participants’ data in the PET analysis, we created a study-specific PET template using Advanced Normalization Tools (ANTs) (25). The template we created was an average of each individual summed PET scan, after mapping onto a common space. We mapped each individual’s FreeSurfer regions-of-interest (ROIs) to this custom template again using ANTs. These ROIs were binarised and summed together before being thresholded in order to include only voxels in which the striatum was present in more than 50% of participants. This custom striatum mask was then warped back into the native PET space for those subjects who did not have MRI scans using the inverse (template-to-native) transformation that was generated using ANTs. All PET ROIs were visually inspected for accuracy.

Once the ROIs were defined in native PET space, we determined [18F]-DOPA uptake 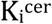 (min^−1^)], for each ROI using the Gjedde-Patlak graphic analysis adapted for a reference tissue input function (26). The cerebellum region was used as the reference region as it represents non-specific uptake (27).

### Statistical analyses

ANOVA was used to test the primary hypotheses that there was an effect of group on whole striatal dopamine synthesis capacity and hippocampal volume. p-values from the ANOVAs were adjusted using FDR correction across striatal subregions (appropriate for positively correlated samples) (28). Additional sensitivity analyses were conducted using an ANCOVA with 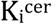 as the dependent variable, group as the independent variable and possible confounds (age, IQ, intra-cranial and striatal ROI volume) as covariates. Separately, in those participants born very preterm, we tested for the independent effects of three neonatal risk factors (perinatal brain injury, gestational age at birth and birth weight) on dopamine synthesis capacity in the whole striatum using an ANCOVA, with 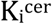 as the independent variable, group (VPT-perinatal brain injury vs VPT-no diagnosed injury) as an independent variable and gestational age at birth and birth weight as covariates. In order to test for regional differences in the effect of VPT birth and perinatal brain injury on dopamine synthesis capacity (analyzing the entire sample), we performed a repeated-measures ANOVA with striatal subregion as the within-subjects factor, group as the between-subjects factor and 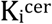 as the dependent variable. To test for a hippocampal volume by striatal subregion interaction, we again performed a repeated-measures ANOVA, with subregion as the within-subject factor, hippocampal volume and intracranial volume as covariates and and 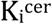 as the dependent variable. To examine whether the relationship between hippocampal volume and striatal 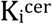 varied significantly by group, we used region as a within-subjects factor, with group and the group-by-hippocampal-volume interaction term as between-subjects factors. A two tailed p value<0.05 was taken as significant.

## Results

### Participants

Seventeen individuals from the VPT-perinatal brain injury group, fourteen from the VPT-no diagnosed injury group and fourteen from the term-born control group were recruited. One VPT-perinatal brain injury participant was excluded from both PET and MRI analysis as a diagnosis of hypothyroidism was discovered at assessment. Incomplete PET data were acquired in one subject from the VPT-no diagnosed injury group because the participant felt unwell and finished the PET scan early. This participant was also excluded from further analysis. In addition to the two participants (one perinatal brain injury, one very preterm no diagnosed injury) excluded from the PET study, four further participants (three perinatal brain injury, one control) were not included in the MRI study due to contraindications to scanning. Thus, thirteen individuals from the VPT-perinatal brain injury group, thirteen from the VPT-no diagnosed injury group and thirteen from the term-born control group had complete PET-and MRI-derived measures.

VPT-perinatal brain injury participants had a lower gestational age and birth weight than VPT-no diagnosed injury participants (Table 1). This was expected as lower gestational age at birth and birth weight are strongly associated with increased risk of perinatal brain injury (29). There were no group differences in age at scanning, IQ, injected dose, gender, alcohol consumption, smoking or socio-economic status between the groups in the PET sample (Table 1).

**Table 1.** Neonatal, socio-demographic, cognitive and scanning measures.

|  | Very preterm-perinatal brain injury<br>(n=16) | Very preterm-no diagnosed injury<br>(n=13) | Controls<br>(n=14) | Test statistic | Significance |
| --- | --- | --- | --- | --- | --- |
| Gestational age in weeks<br>Mean (SD) | 28.44<br>(2.28) | 30.46<br>(1.76) | | $U_{27} = 47.00$ | $p = 0.011$ |
| Birth weight in grams<br>Mean (SD) | 1203.19<br>(304.95) | 1557.15<br>(364.98) | | $U_{27} = 46.50$ | $p = 0.012$ |
| Age in years<br>Mean (SD) | 30.21<br>(1.78) | 30.85<br>(2.09) | 29.81<br>(3.24) | $F_{2,40} = 1.50$ | $p = 0.236$ |
| Sex<br>(female:male) | 03:13 | 04:09 | 05:09 | $X^2_2 = 1.14$ | $p = 0.564$ |
| High SES (%)* | 68.75 | 69.23 | 61.53 | $X^2_2 = 0.22$ | $p = 0.894$ |
| IQ<br>Mean (SD) | 106.67<br>(14.52) | 107.73<br>(10.07) | 110.40<br>(10.52) | $F_{2,33} = 0.28$ | $p = 0.755$ |
| Alcohol consumption<br>(Units/week) | 7.40<br>(11.30) | 12.50<br>(11.99) | 5.50 (4.72) | $X^2_2 = 3.172$ | $p = 0.205$ |
| Injected dose (MBq)<br>Mean (SD) | 146.44<br>(2.15) | 146.25<br>(2.52) | 145.73<br>(2.38) | $F_{2,40} = 0.23$ | $p = 0.793$ |

\*SES was collapsed into two groups; the percent of participants belonging to the high SES (level 1–2) category is presented in the table.

### Dopamine synthesis capacity

There was a significant effect of group on 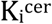 corresponding to a partial eta-squared of 0.233 (a large effect size, Table 2). Post-hoc tests showed 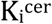 was significantly reduced in the VPT-perinatal brain injury group compared to the VPT-no diagnosed injury group (p = 0.023, Cohen’s d = 1.36) and controls (p = 0.010, Cohen’s d = 1.07) in the whole striatum with large effect sizes (Figure 1, Table 2). There was no significant difference in 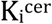 between the VPT-no diagnosed injury group and controls (Figure 1, Table 2).

**Figure 1.**
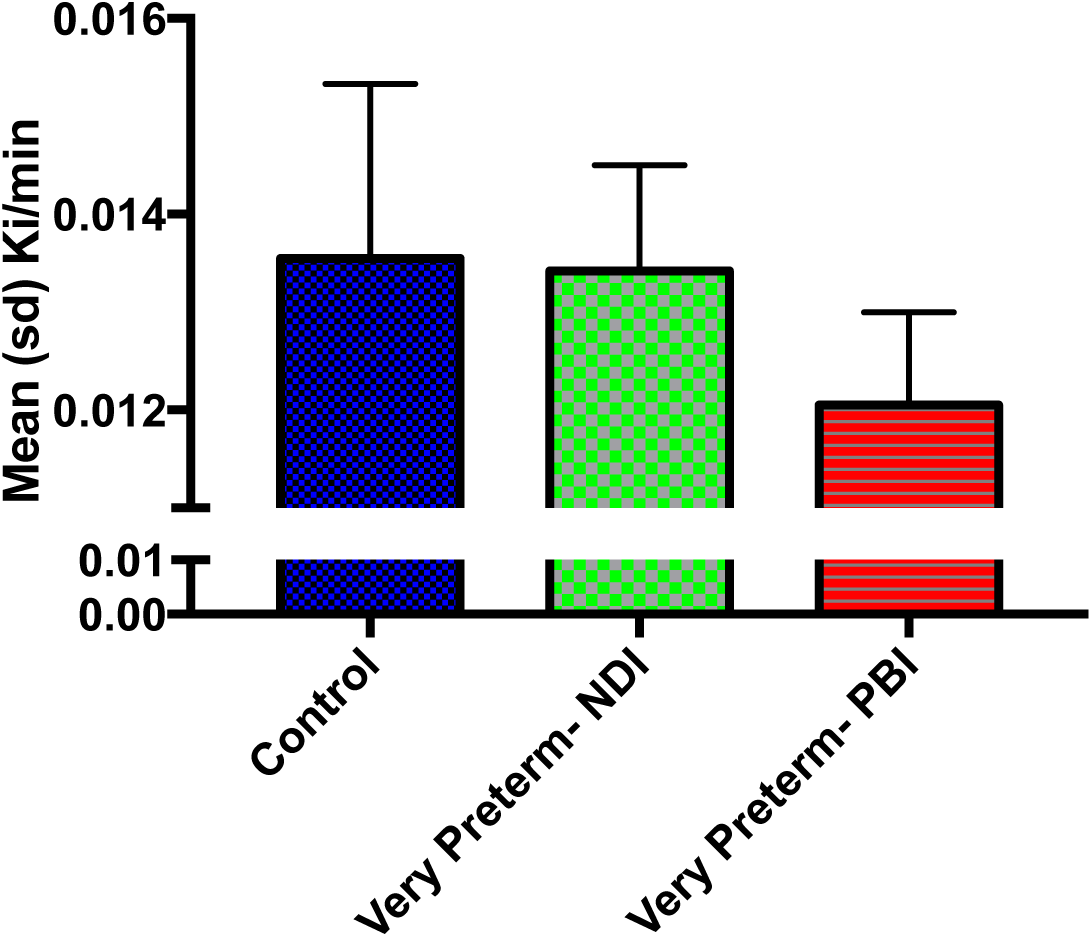
Whole striatal dopamine synthesis capacity by group

Individuals who suffered macroscopic perinatal brain injury related to VPT birth (VPT-perinatal brain injury) had significantly lower dopamine synthesis capacity in the whole striatum compared to other adults born VPT with no macroscopic perinatal brain injury (corrected p = 0.023, Cohen’s d = 1.36) and full term-born controls (corrected p = 0.01, Cohen’s d = 1.07).

The reduction in dopamine synthesis capacity was significant in the caudate nucleus and the nucleus accumbens, but not the putamen (see Table 2).

**Table 2.**
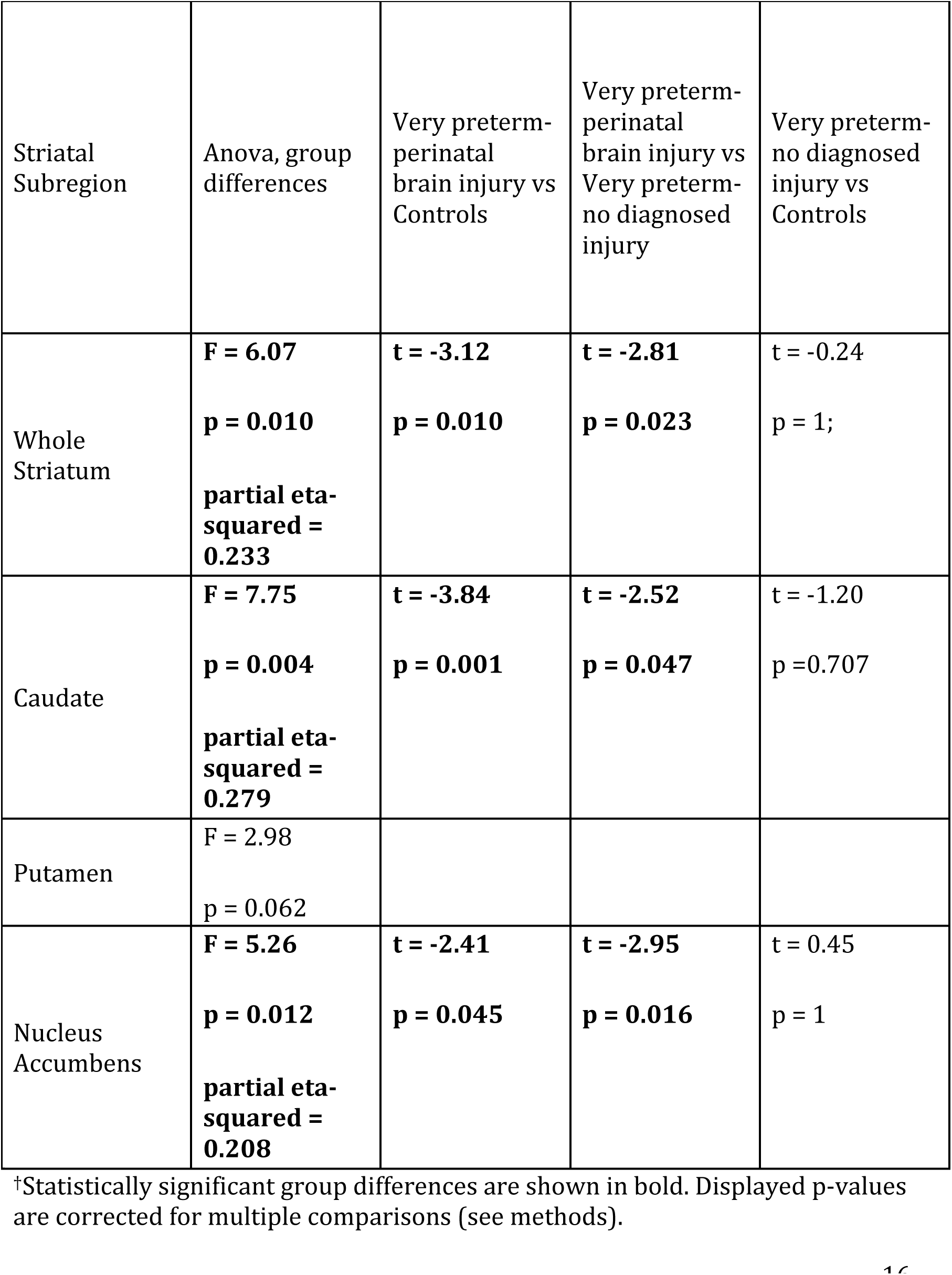
Striatal subregion dopamine synthesis capacity†.

| Striatal Subregion | Anova, group differences | Very preterm-perinatal brain injury vs Controls | Very preterm-perinatal brain injury vs Very preterm-no diagnosed injury | Very preterm-no diagnosed injury vs Controls |
| --- | --- | --- | --- | --- |
| Whole Striatum | **F = 6.07**<br>**p = 0.010**<br>**partial eta-squared = 0.233** | **t = -3.12**<br>**p = 0.010** | **t = -2.81**<br>**p = 0.023** | t = -0.24<br>p = 1; |
| Caudate | **F = 7.75**<br>**p = 0.004**<br>**partial eta-squared = 0.279** | **t = -3.84**<br>**p = 0.001** | **t = -2.52**<br>**p = 0.047** | t = -1.20<br>p = 0.707 |
| Putamen | F = 2.98<br>p = 0.062 |  |  |  |
| Nucleus Accumbens | **F = 5.26**<br>**p = 0.012**<br>**partial eta-squared = 0.208** | **t = -2.41**<br>**p = 0.045** | **t = -2.95**<br>**p = 0.016** | t = 0.45<br>p = 1 |

†Statistically significant group differences are shown in bold. Displayed p-values are corrected for multiple comparisons (see methods).

Additional sensitivity analyses showed that the reduction in 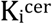 in the VPT-perinatal brain injury group remained significant when removing all participants who had a history of psychiatric diagnosis (VPT-perinatal brain injury group n=4, VPT-no diagnosed injury group n=2, control group n=1) in the whole striatum (F = 4.825, p = 0.023) and the caudate nucleus (F = 5.608, p = 0.023) but not the nucleus accumbens (F = 3.047, p = 0.061). Furthermore, when just including the participants who also took part in the MRI study (and hence had individual FreeSurfer-based striatal segmentations), reduced 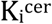 in the VPT-perinatal brain injury group remained significant in the whole striatum (F = 5.708, p = 0.018), the caudate nucleus (F = 10.130, p = 0.003) and in the nucleus accumbens (F = 4.306, p = 0.034).

The reduction in 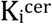 in the VPT-perinatal brain injury group remained significant when co-varying for age, IQ, region-of-interest (i.e. whole striatum, caudate, putamen or nucleus accumbens) volume and intracranial volume in the whole striatum (F = 7.113, p = 0.005), the caudate nucleus (F = 7.083, p = 0.005) and in the nucleus accumbens (F = 3.663, p = 0.037).

Our VPT groups differed not only on perinatal brain injury status, but also on gestational age at birth and birth weight (Table 1). Furthermore, younger gestational age and lower birth weight are both common risk factors for perinatal brain injury (29). When we combined these three neonatal risk factors into a single model to predict whole striatal 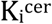, perinatal brain injury remained a significant predictor of dopamine synthesis capacity (F = 9.23, p = 0.006), but neither gestational age at birth (F = 0.01, p = 0.929), nor birth weight (F = 0.01, p = 0.925) significantly predicted dopamine synthesis capacity.

In order to further probe whether group differences in 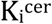 varied across striatal subregions we performed a repeated-measures ANOVA with striatal subregion as the within-subjects factor, group as the between subjects factor and 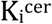 as the dependent variable. There was no significant subregion-by-group interaction (F = 1.03, p = 0.398). As expected there were significant effects of subregion (F = 81.26 p < 0.001) and group (F = 6.95, p = 0.003).

### Hippocampal and striatal volume analysis

There was a significant difference in hippocampal volumes across the three groups (Table 3). The VPT-perinatal brain injury group had significantly lower volumes than controls, while the VPT-no diagnosed injury group did not differ significantly from either group (Table 3). The group differences in hippocampal volume remained significant after controlling for intracranial volume (ICV) (F = 7.19, p = 0.002).

On assessing striatal volume with repeated-measures ANOVA, with striatal subregion volume as a within-subjects factor and group as a between-subjects factor, we found no significant main effect of group (p = 0.081) and no significant group*subregion interaction (p = 0.123).

Analysing the whole striatum and each sub-region separately using one-way ANOVAs and post-hoc t-tests confirmed that there were no significant between-group volumetric differences in the striatum (Table 3).

We additionally analysed the estimated striatal volumes for all individuals with PET scans (i.e. including those without MRI), and again found that there were no significant between group differences in striatal volume as a whole (F = 0.77, p = 0.628) or in any striatal subregion after FDR correction for multiple comparisons (caudate, F = 0.17, p = 0.841; putamen, F = 2.73, p = 0.154), although there was a trend for differences between groups in the volume of the nucleus accumbens, which did not reach significance (F = 4.38, p = 0.076).

**Table 3.** Subcortical volumes (mm3) ‡

|  | Very preterm-perinatal brain injury<br>mean (sd) | Very preterm – no diagnosed injury<br>mean (sd) | Control<br>mean (sd) | Anova | Very preterm-perinatal brain injury vs Control | Very preterm-perinatal brain injury vs very preterm-no diagnosed injury | Very preterm-no diagnosed injury vs Control |
| --- | --- | --- | --- | --- | --- | --- | --- |
| Hippocampus | 8624 (1329) | 9557 (1113) | 10090 (1329) | **F = 4.928**<br>**p = 0.013** | **t = -3.10**<br>**p = 0.011** | t = -1.97<br>p = 0.168 | t = -1.13<br>p = 0.802 |
| Striatum (whole) | 19098 (3217) | 19767 (2495) | 21487 (2322) | F = 2.70<br>p = 0.081 | t = -2.17<br>p = 0.092 | t = -0.59<br>p = 1 | t = -1.82<br>p = 0.341 |
| Caudate | 7491 (1516) | 7609 (998) | 8250 (910) | F = 1.581<br>p = 0.22 | t = -1.55<br>p = 0.322 | t = -0.23<br>p = 1 | t = -1.71<br>p = 0.514 |
| Putamen | 10541 (1682) | 10973 (1480) | 11977 (1655) | F = 2.824<br>p = 0.11 | t = -2.25<br>p = 0.079 | t = -0.70<br>p = 1 | t = -1.68<br>p = 0.343 |
| Nucleus Accumbens | 1066 (210) | 1185 (167) | 1260 (227) | F = 3.023<br>p = 0.11 | t = -2.26<br>p = 0.06 | t = -1.60<br>p = 0.43 | t = -0.96<br>p = 1 |

‡Statistically significant group differences are shown in bold. p-values for ANOVA
tests of the striatal subregions adjusted using FDR method for positively
correlated samples. p-values for the post-hoc t-tests are corrected for multiple
comparisons using the Bonferroni method.

### Dopamine synthesis capacity and hippocampal volume

A significant correlation was observed between hippocampal volume and 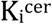 in the caudate (r = 0.34, p = 0.032, Figure 2A) and in the nucleus accumbens (r = 0.32, p = 0.049, Figure 2B) across the whole sample. These associations remained significant when controlling for ICV (caudate 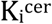-hippocampal volume, r = 0.39, p = 0.017; nucleus accumbens 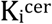, r = 0.34, p = 0.036).

In order to test the interaction between hippocampal volume and striatal subregion, we again performed a repeated-measures ANOVA, with subregion as the within-subject factor, hippocampal volume and intracranial volume as covariates and and 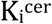 as the dependent variable. We found a significant effect of hippocampal volume (F = 4.90, p = 0.033), but no hippocampal volume by striatal subregion interaction (F = 0.88, p = 0.420). Additionally, there was no significant effect of ICV on 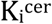 (F = 1.11, p = 0.299). We then examined whether the relationship between hippocampal volume and striatal 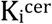 varied significantly by group, again by using region as a within-subjects factor, and this time having group and the group-by-hippocampal-volume interaction term as between-subjects factors. Again we found a significant main effect of group (F = 4.794, p = 0.015), but no group by hippocampal volume interaction (F = 0.41, p = 0.747).

**Figure 2.**
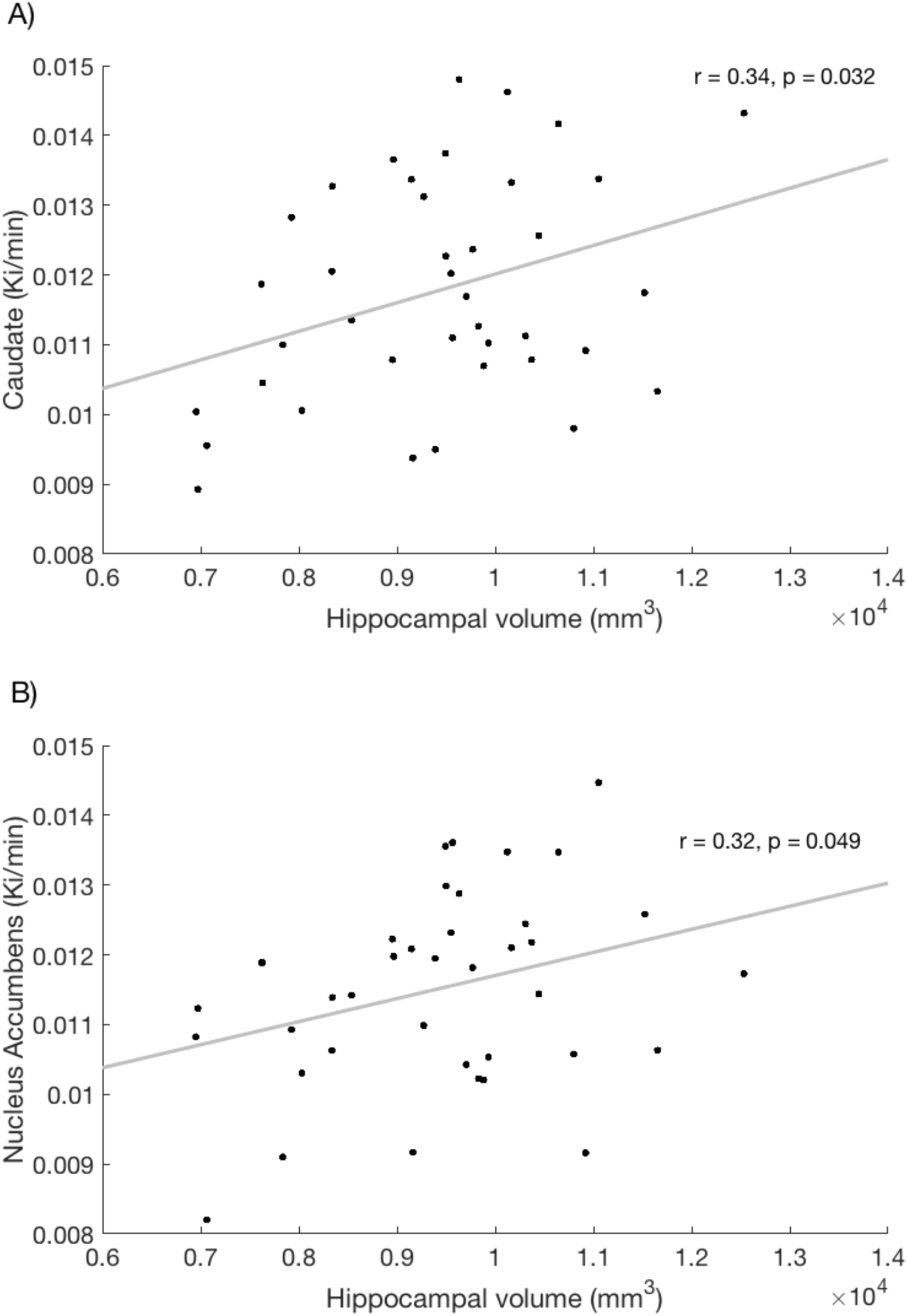
Relationship between hippocampal volume and dopamine synthesis capacity in the A) caudate and B) nucleus accumbens.

## Discussion

### Main findings

Adults with a history of macroscopic perinatal brain injury associated with VPT birth had reduced dopamine synthesis capacity in the striatum compared to controls born VPT and those born at term, and reduced hippocampal volume compared to individuals born at term. Individuals born similarly preterm but without evidence of macroscopic brain injury showed no significant differences in presynaptic dopamine synthesis capacity from controls, suggesting that preterm birth in the absence of macroscopic brain injury is not sufficient to disrupt striatal dopaminergic function in adult life.

### Possible Mechanisms

It is possible that perinatal brain insults resulted in a long-lasting reduction in the number of dopaminergic neurons (30, 31) or caused a down-regulation in dopamine synthetic enzyme levels, in line with post-mortem findings showing reduced tyrosine hydroxylase expression in dopaminergic neurons following prolonged hypoxia (12). One alternative possibility is that a common genetic or environmental cause predisposes to both low striatal dopamine synthesis and the direct causes of perinatal brain injury.

### Hippocampus and striatal dopamine

We also found that reduced striatal dopamine synthesis capacity was associated with reduced hippocampal volume. Preclinical models have shown that hippocampal damage can result in altered striatal dopaminergic function (32–34). However, dopamine also has effects on neurodevelopment, influencing neuronal migration, neurite outgrowth and synapse formation (35), and these effects are particularly marked during the second half of a typical pregnancy (36), indicating that dopaminergic changes could also influence hippocampal development. Untangling the timing of dopaminergic or hippocampal alterations would seemingly require serial measurements of both systems over the perinatal period, which likely requires post-mortem or preclinical studies.

### Cognitive Implications

These results may have implications for cognitive function in people born preterm. While the current group of study participants were not cognitively impaired, cognitive deficits are commonly found in individuals born VPT, and are exacerbated following perinatal brain injury (37). Both longitudinal studies of individuals born preterm and preclinical studies have suggested a link between neonatal hippocampal injury and later working memory impairments (7, 38, 39). Our findings are consistent with preclinical work suggesting a link between early hippocampal injury and altered development of the dopaminergic system (40), which is crucial for cognitive functions such as reward-based learning (41) and working memory (42). Working memory is a particularly common deficit in children born with perinatal brain injury (43, 44), and is associated with academic outcome in this population (45). Individuals with lower presynaptic dopamine synthesis in the caudate nucleus tend to have worse working memory performance (46, 47) and respond better to dopamine agonists as cognitive enhancers than individuals with higher baseline dopamine synthesis (48). Our findings suggest that VPT individuals with perinatal brain injury who experience cognitive difficulties could benefit from dopamine agonists as cognitive enhancers, perhaps by dopamine’s role in enhancing intrinsic plasticity mechanisms (49) that have been observed in this population (50).

### Relationship with Psychiatric Disorders

Reduced dopamine synthesis capacity is also associated with substance dependence (51, 52), major depression (53) and Parkinson’s disease (54). Our findings thus suggest that people with perinatal brain injury could be at increased risk for a number of neuropsychiatric disorders.

### Obstetric Complications and Schizophrenia

In contrast, dopamine synthesis capacity is increased in the majority of people with schizophrenia (13) and people at risk of schizophrenia (55). As yet there have been no PET studies specifically of those people with schizophrenia who have had severe obstetric complications, although it is known that they are especially likely to have small left hippocampi (56). Nevertheless, it is not clear how our results fit with findings that obstetric complications increase the risk of schizophrenia, where interaction with genetic risk factors is likely to be involved (57, 58).

#### Mechanism linking Perinatal Brain Injury with Psychosis Risk

In contrast to the increased dopamine synthesis capacity seen in most schizophrenia patients, those who develop schizophrenia-like psychoses following abuse of drugs (59), and those with treatment resistant schizophrenia do not share this increased synthesis capacity (60). It is thus possible that the relationship between VPT birth, perinatal brain injury and increased risk for psychosis does not depend on presynaptic dopamine synthesis capacity. It may be important to closely monitor the condition of those individuals born VPT with perinatal brain injury who are treated with antipsychotic medication, as reducing an already-reduced dopaminergic system could lead to unintended extrapyramidal and cognitive effects. Alternatively, it is possible that hypersensitive postsynaptic dopaminergic D2 receptors could unite the seemingly discordant findings of reduced presynaptic dopamine synthesis and increased psychosis risk, as appears to be the case in substance-dependent patients with schizophrenia (59). If such disruption were to occur during development, it could have dramatic effects on the developing brain (61), with pre-frontal dependent cognitive functions such as working memory perhaps particularly vulnerable (62).

#### Implications for People born VPT without Macroscopic Perinatal Brain Injury

Our finding that there are not marked alterations in dopamine synthesis capacity in the VPT-no diagnosed injury group is also important for the large numbers of people born preterm, as it indicates that the development of the dopamine system, or at least those aspects related to dopamine synthesis, is not disrupted long-term in the absence of macroscopic perinatal brain injury. The VPT-perinatal brain injury and VPT-no diagnosed injury groups in the present study also differed in gestational age, and birth weight, as these neonatal risk factors tend to co-occur (29). Nonetheless, when all three factors were introduced in the same model, only perinatal brain injury was a significant predictor of adult dopamine synthesis capacity. This suggests that reduced striatal dopamine synthesis capacity in adulthood is specific to those individuals with perinatal brain injury.

### Limitations

From a methodological perspective it is possible that between-group differences in the accuracy of image registration may contribute to the apparent reduction in dopamine synthesis capacity seen in the VPT-perinatal brain injury group. However, we used the subject’s own MRI to define the PET region of interest which should mitigate, although not entirely avoid, this risk. Moreover, the results remained significant after controlling for both striatal and total intracranial volume or excluding subjects without MRI scans, suggesting that volume reductions or normalisation differences do not account for the findings. The postnatal ultrasound scans exclude macroscopic brain injury in the VPT-no diagnosed injury group but do not exclude a variety of other microscopic alterations. However, this would not explain our results, as it would, if anything, reduce group differences.

### Conclusions

In summary, we found reduced presynaptic dopamine synthesis capacity in the striatum in individuals born VPT with macroscopic perinatal brain injury. This may help to guide pharmacological interventions for cognitive deficits in this group. We additionally found significant associations between dopaminergic function and reduced hippocampal volume. These results indicate there are longterm neurochemical and structural consequences of perinatal brain injury.

Supporting data is available on request: please contact

## Acknowledgements

We would like to thank our participants and funders. This study was funded by the March of Dimes (grant number #12-FY11-206) and Medical Research Council-UK (MRC MR/K004867/1) grants to Dr. Nosarti and Medical Research Council-UK (no. MC-A656-5QD30), Maudsley Charity (no. 666), Brain and Behavior Research Foundation, and Wellcome Trust (no. 094849/Z/10/Z) grants to Dr. Howes and the National Institute for Health Research (NIHR) Biomedical Research Centre at South London and Maudsley NHS Foundation Trust and King’s College London. The views expressed are those of the author(s) and not necessarily those of the NHS, the NIHR or the Department of Health.

